# Comparative effects of oncogenic mutations G12C, G12V, G13D, and Q61H on local conformations and dynamics of K-Ras

**DOI:** 10.1101/2020.03.26.010710

**Authors:** Sezen Vatansever, Burak Erman, Zeynep H. Gümüş

## Abstract

K-Ras is the most frequently mutated protein in human cancers. However, until very recently, its oncogenic mutants were viewed as undruggable. To develop inhibitors that directly target oncogenic K-Ras mutants, we need to understand both their mutant-specific and pan-mutant dynamics and conformations. Recently, we have investigated how the most frequently observed K-Ras mutation in cancer patients, G12D, changes its local dynamics and conformations^1^. Here, we extend our analysis to study and compare the local effects of other frequently observed oncogenic mutations, G12C, G12V, G13D and Q61H. For this purpose, we have performed Molecular Dynamics (MD) simulations of each mutant when active (GTP-bound) and inactive (GDP-bound), analyzed their trajectories, and compared how each mutant changes local residue conformations, inter-protein distance distributions, local flexibility and residue pair correlated motions. Our results reveal that in the four active oncogenic mutants we have studied, the α2 helix moves closer to the C-terminal of the α3 helix. However, P-loop mutations cause α3 helix to move away from Loop7, and only G12 mutations change the local conformational state populations of the protein. Furthermore, the motions of coupled residues are mutant-specific: G12 mutations lead to new negative correlations between residue motions, while Q61H destroys them. Overall, our findings on the local conformational states and protein dynamics of oncogenic K-Ras mutants can provide insights for both mutant-selective and pan-mutant targeted inhibition efforts.

## 1. INTRODUCTION

K-Ras is a small GTPase that plays a crucial role in cellular signaling and promotes cellular proliferation, survival, growth and differentiation^2^. The protein controls signaling networks by functioning as a molecular switch that cycles between an inactive GDP-bound and an active GTP-bound state^3,4^. The balance between the two states is regulated by guanine nucleotide exchange factors (GEFs) that bind to inactive K-Ras (K-Ras-GDP) and stimulate the exchange of GDP with GTP. After GTP binding, K-Ras becomes active (K-Ras-GTP) and can bind and activate its downstream effector proteins, such as Raf kinase, phosphatidylinositol 3-kinase (PI3K), and Ral guanine nucleotide dissociation stimulator (RalGDS) ^5-7^. To terminate the downstream signaling, active K-Ras catalyzes GTP hydrolysis to return to its inactive state. The intrinsic GTPase activity of K-Ras-GTP can be enhanced by GTPase-activating protein (GAP) binding^8,9^. A complete GTPase reaction requires well-ordered conformations of the protein active site, which includes the P-loop (residues 10-17), switch I (SI, residues 25-40) and switch II (SII, residues 60-74) regions (Fig 1).

**Figure 1.**
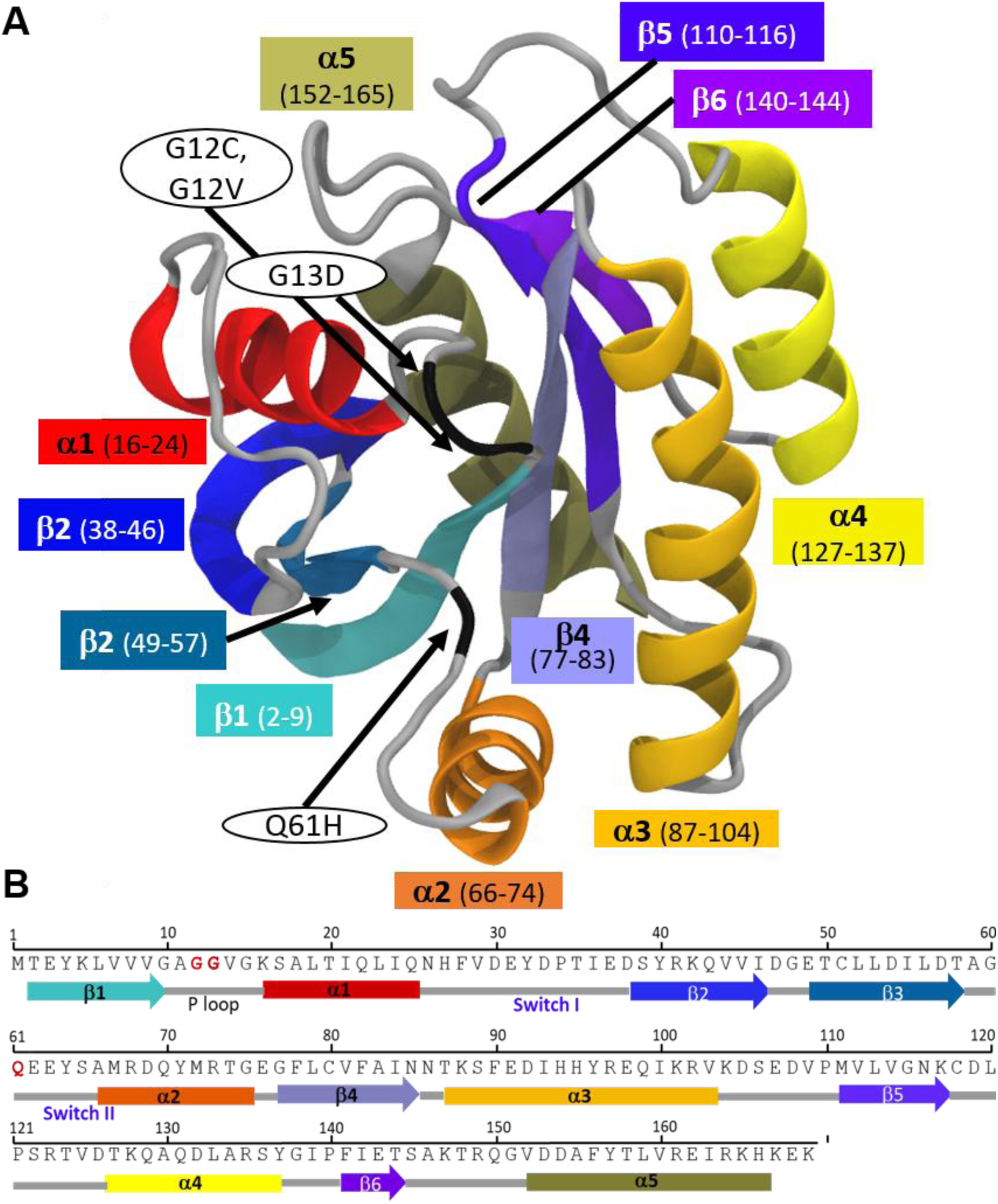
K-Ras protein and the most frequently mutated residues. (A) Secondary structure of K-Ras in ribbon representation. Functional regions are in the same color as in K-Ras sequence. Arrows point to mutated residues. (B) Schematic of K-Ras sequence (residues 1–169). Arrows: β-sheets, rectangles: α-helices.

Based on The Catalog of Somatic Mutations in Cancer (COSMIC), the most frequently observed oncogenic K-Ras mutations in cancer patients are at active site residues G12 (89%), G13 (9%) and Q61 (1%)^5,10^. These mutations impair both intrinsic and GAP-accelerated GTPase activity while increasing the nucleotide exchange activity, which disturbs the balance between active and inactive states^11,12^. Since mutant K-Ras-GTP cannot return back to its inactive form, it continuously triggers the downstream signaling networks that are ultimately related with oncogenic cellular growth ^5,13-15^. However, blocking the continuous activity of oncogenic K-Ras with selective mutant or pan-mutant inhibition remains a formidable task^16,17^.

Compelling evidence suggest that there are distinct mutation-specific effects on downstream effector signaling pathways^18^. Ihle et al. showed mutation-specific changes in downstream pathways in non-small cell lung cancer (NSCLC) cell lines^19^. Specifically, they observed that while both G12C and G12D mutations activate Ral signaling and decrease growth factor-dependent Akt activation, only G12D mutation activates phosphatidylinositol 3-kinase (PI-3-K) and mitogen-activated protein/extracellular signal-regulated kinase (MEK) signaling. In another study, Hammond *et al*. investigated the distinct effects of several KRAS mutations (G12V, G12D and G13D) on KRAS-mediated pathways by using quantitative analysis of the proteome of isogenic SW48 colon cancer cell lines^20^. They found that while G12 mutations induce the colon cancer stem cell marker DCLK1 and the receptor tyrosine kinase, the G13D mutation induces the tight-junction protein ZO-2. These studies suggest that K-Ras mutants can have distinct effects on downstream signaling pathways in cancer.

While distinct effects of different K-Ras mutations have been described in some studies, most studies so far have treated all K-Ras mutants as a single entity, considering the protein as either wild-type or mutant ^21^. To understand the conformational and dynamic changes due to the mutations, researchers have utilized molecular dynamics (MD) simulations and described the distinct global dynamics of mutant complexes (i.e., mutant K-Ras proteins in complex with GTP/GDP)^22,23^. However, identifying the unique local dynamics of K-Ras specific to its oncogenic mutation can also help understand the individual characteristics of each mutant protein, which can assist in the development of targeted therapies. In our previous work, we have presented a detailed study on the effects of the most recurrent oncogenic mutation, G12D, on local dynamic characteristics of both active and inactive K-Ras using long timescale MD simulations, and observed nucleotide-specific effects on local conformations and dynamics of the protein.

Following up on our previous study on the G12D mutant, here, we present mutation-specific (and agnostic) effects of other frequently observed oncogenic K-Ras mutants, including G12C, G12V, G13D and Q61H on local protein conformations and dynamics and provide an atomistic-level explanation for these effects. For this purpose, we have performed long timescale MD simulations of each mutant in both GTP-bound active and GDP-bound inactive forms and compared them with each other and the wild-type protein. Briefly, we have first identified the individual effects of each mutation on local residue conformations by calculating the changes in intra-protein residue pair distances. Then, we have identified the favored conformations of residue pairs in each mutant protein complex by plotting the distributions of their distances. These provided information on how each oncogenic mutation alters the local conformational dynamics of active and inactive wild-type K-Ras. We next asked whether these oncogenic mutations caused certain protein regions to become more flexible or rigid. To understand and quantify mutation-specific local changes in protein flexibility, we compared the residue fluctuations of each mutant in both active and inactive form with those of wild-type K-Ras. Next, we aimed to understand the regulation of local protein dynamics by the allosteric coupling of residue fluctuations in each mutant system. For this purpose, we described the regulation of local protein motions by plotting the pair-wise correlations of residue fluctuations. Each mutant displayed distinct patterns in residue-residue correlation maps that revealed mutation-specific regulation of local dynamics. In summary, we have analyzed and compared the local dynamics of each oncogenic K-Ras mutant in both active and inactive forms with the wild-type protein, which revealed mutation-specific effects on local protein conformations and dynamics. We anticipate that our results will inform future studies on selective targeting of K-Ras oncogenic mutants in their active or inactive states.

## 2. RESULTS

### 2.1. CONFORMATIONAL CHANGES IN K-RAS DUE TO ONCOGENIC MUTATIONS G12C, G12V, G13D AND Q61H

#### 2.1.1 In active K-Ras, residue G12 mutations cause SII to move away from both the α3 helix and the P-loop

To explore how oncogenic mutations alter the local residue conformations of K-Ras, we first performed MD simulations of the GTP- and GDP-bound oncogenic K-Ras mutants G12C, G12V, G13D and Q61H. Next, we analyzed each trajectory using a protocol from our previous study on G12D^1^, which we summarize in Methods. Briefly, for each GTP-bound protein, to understand the contribution of non-bonded residue-residue pairs to changes in local conformation^29,30^, we defined a sphere with radius ∼ 9.1 Å around each residue, and calculated the time-averaged distance 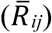 of all residues to the central residue within this sphere. Such a sphere around a residue includes both its contacting neighbors and non-bonded local interactions, and corresponds to its *second coordination shell*, as conceptualized within the popularly used protein dynamics analysis approach of Gaussian Network Modeling (GNM)^24-27^ (see Methods). We then quantified the changes in residue-residue distances 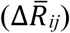 caused by each mutation by assuming K-Ras^WT^ as the reference structure and used the first frame of its trajectory to determine the residue pairs (*ij*) within the second coordination shell. In Figure 2, we show the 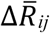 values for all residue pairs (*ij*), where positive values correspond to regions that move away from each other upon mutation and negative values correspond to regions that get closer. As suggested by the 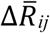 values in Figures 2A and 2C, the SII loop (Q61-E62) moves away from the α3 helix (D92-H95) in G12 mutants G12C and G12V. Furthermore, SII residue Q61 also moves away from the P-loop (A11-G13).

**Figure 2.**
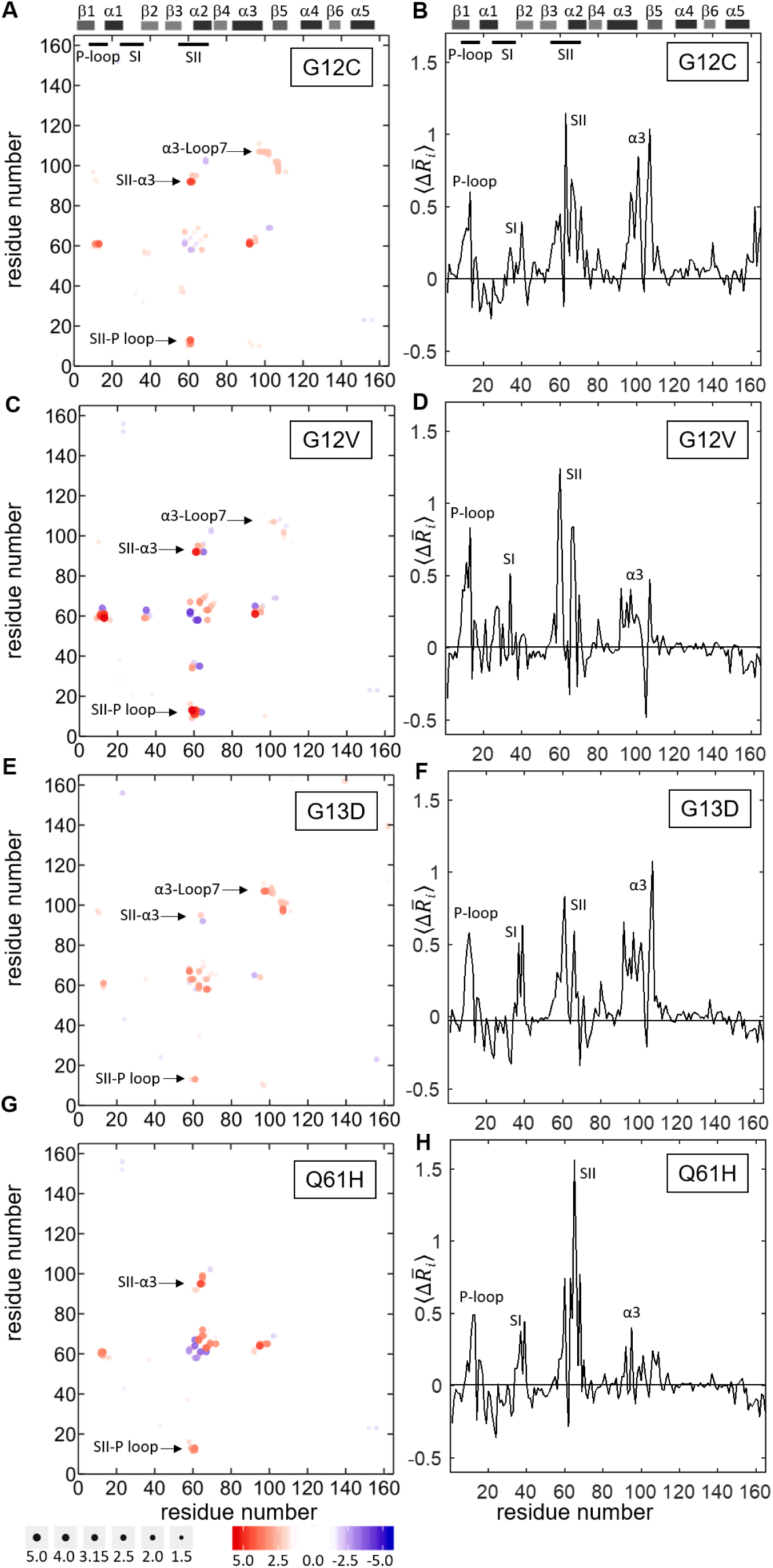
Alterations in active K-Ras conformations due to oncogenic mutations. Left panels: changes in pairwise residue distances 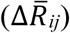 in active K-Ras due to mutation. Positive 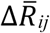 values indicate divergent pairs (red); negative 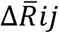 values indicate convergent pairs (blue). Right panels: all 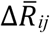 values averaged for each residue, 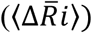. Positive values indicate that the mutation causes a residue to move away from its neighbors; negative values indicate that a residue moves close to its neighbors. The predominant behavior for all studied mutants is positive (A-B) K-Ras^G12C^ (C-D) K-Ras^G12V^ (E-F) K-Ras^G13D^ (G-H) K-Ras^Q61H^.

MD simulations of the mutant K-Ras-GTP complexes revealed that similar to our observations in the G12D mutant in our earlier study^1^, the salt bridge between E62 (SII) and K88 (α3) in K-Ras^WT^-GTP disappears in other G12 mutants. Combined with our observations in Figure 2, these results suggest that SII moves away from its neighbors as a result of the disruption of the salt bridge between SII and α3. Intriguingly, in G13D mutant, the deviation of SII from its neighbors is not as remarkable as in G12 mutants.

In addition to the changes in SII distances, we also observed that the α3 helix (Q99-K101) moves away from the Loop7 (E107) in G12C, G12V and G13D mutants (but not Q61H), as shown in Figure 2.

#### 2.1.2. In active K-Ras, studied oncogenic mutations cause the P-loop, SI, SII and α3 regions to move away from their neighbors

In addition to analyzing the mutation effects on pairwise distances 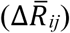, we also quantified the effects on individual conformations of residues relative to their neighbors’. Specifically, for each oncogenic mutant, we separately plot the average pairwise distance of each residue, 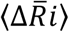, as shown in Figure 2. These plots revealed that in all oncogenic K-Ras mutants, the P-loop, SI, SII and the α3 helix regions become distant from their neighbors, with the effect most pronounced in G12C and G13D mutants. However, the G12V mutant exhibits large deviations in both P-loop and SII, while the Q61H mutant shows distinctly strong deviations in SII.

In summary, based on our time-averaged inter-residue distance analysis, oncogenic mutations studied cause largest local conformational changes in the SII, α3 and P-loop regions (Fig 2). These analyses provide static information on mutational effects on local conformations. Next, to gain dynamic information, we investigated how the local conformations distributed over the simulation time, as described below.

#### 2.1.3. Oncogenic mutations at residue G12 alter the balance of residue pair distances between SII and α3 helix

To better understand the dynamic effects of mutations on local residue conformations in the regions that show the largest static deviations from the wild-type, we plot the distance distributions between Cα-Cα atoms of residue pairs in these regions for each mutant protein in both active and inactive form (Fig 3-5). For a given residue pair, these distribution plots reveal whether the residue pair exhibits: (i) similar conformations in both wild-type and mutant protein simulations, leading to overlapping distribution patterns; or (ii) different conformations in the mutant than in the wild-type, leading to dissimilar distribution patterns. In addition, multiple peaks suggest that the residue pair obtains multiple conformations during the simulation, while a single peak suggests that the residue pair gets stuck in a single conformation.

**Fig 3.**
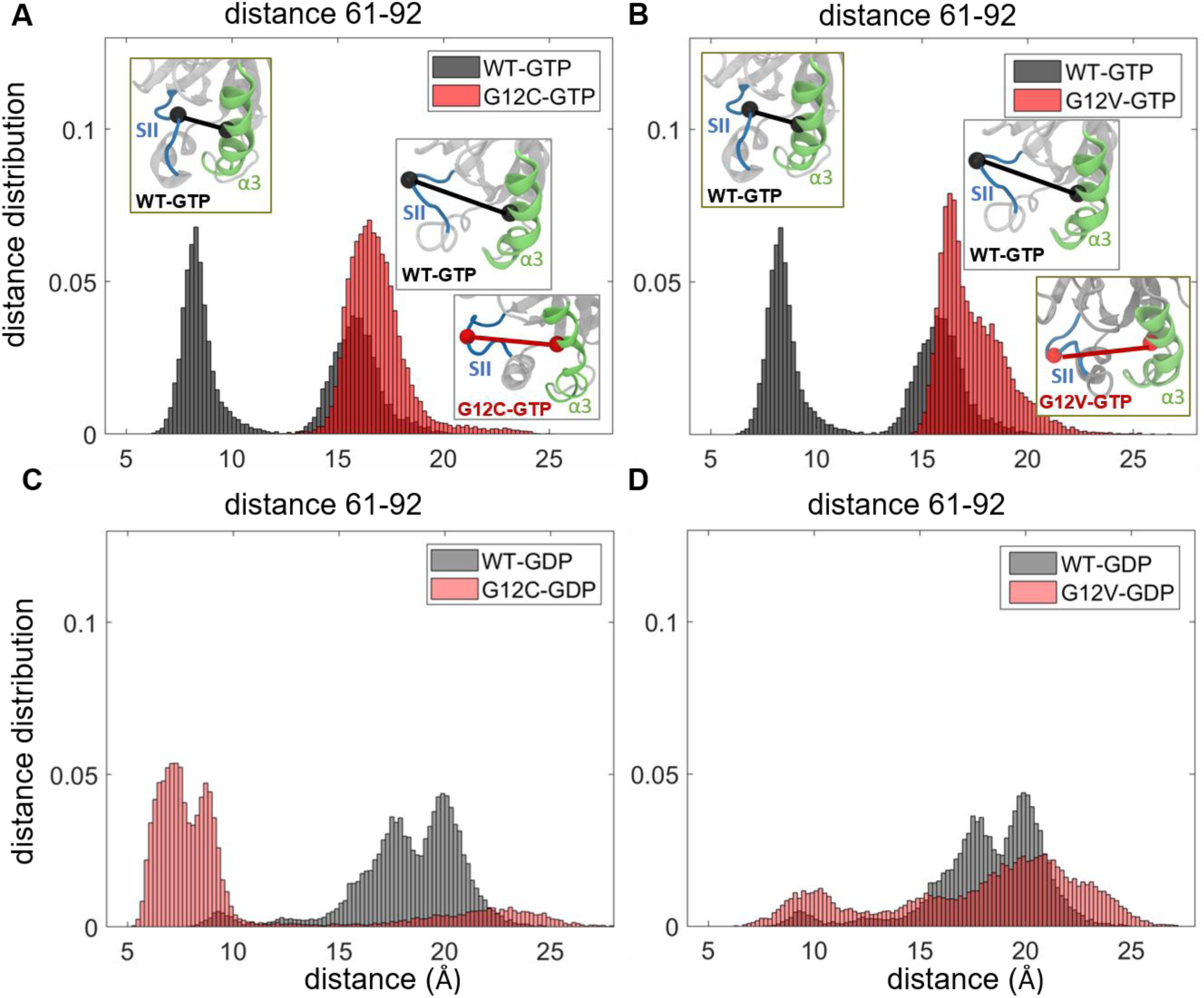
Distribution of distances *W*(*R*_*ij*_) between Cα-Cα atoms of residue pairs in SII-α3 region in wild-type and G12 mutant K-Ras proteins. GTP-bound active K-Ras (black: WT, red: mutant) (A) Q61-D92 in WT and G12C, (B) Q61-D92 in WT and G12V; GTP-bound active K-Ras (grey: WT, pink: mutant) (C) Q61-D92 in WT and G12C, (D) Q61-D92 in WT and G12V.

From these plots, we observed that the distance distribution patterns exhibit distinct characteristics in each mutant system, suggesting that the oncogenic mutations we studied change the balance of distances between protein regions. Specifically, in *active K-Ras*^*WT*^, the distance distribution plots of SII loop-α3 region show only two peaks with narrow distributions (Fig 3A-B, S1A-B). The first peak is at ∼8.5 Å for Q61-92 (8 Å for 62-92) and the second peak is at ∼16.5 Å for Q61-92 (∼13 Å for 62-92), suggesting both close and distant conformations of the residue pairs, respectively. However, both C and V mutations at G12 decrease the number of conformations, leading to distribution curves with only a single peak at ∼16.5 Å for Q61-92 (∼13 Å for 62-92) (Figs 3A-B and S1A-B). These results suggest that after both G12C and G12V mutations, the residue pairs in the SII loop-α3 region populate at the distant conformation, which is consistent with the same population shift we observed after G12D mutation in our earlier study^1^.

In *inactive K-Ras*, while the distances in the SII loop-α3 region were more widely spread, with peaks at larger distances in the wild-type inactive protein (Fig 3C-D), the peaks correspond to shorter distances in G12C mutant (Fig 3C, S1C). These results suggest that in inactive K-Ras, the SII loop-α3 region exhibits various conformations in wild-type but assumes a closer conformation after the G12C mutation, which is similar to that of the G12D mutation^1^. However, G12V mutation does not have such an effect, as shown in Figs 3D and S1C.

#### 2.1.4. In both active and inactive K-Ras, mutations on P-loop and SII alter their conformation populations

In *inactive K-Ras*^*WT*^, the residue pair distances between the P-loop residues (A11-G13) and SII residues (Q61-E63) exhibit multimodal distributions related to multiple conformations of this region. However, the distance distribution plots (Fig 4, S2-3) show that the mutations at the P-loop (G12C, G12D and G13D) and SII (Q61H) decrease the number of conformations of this region in inactive protein.

**Figure 4.**
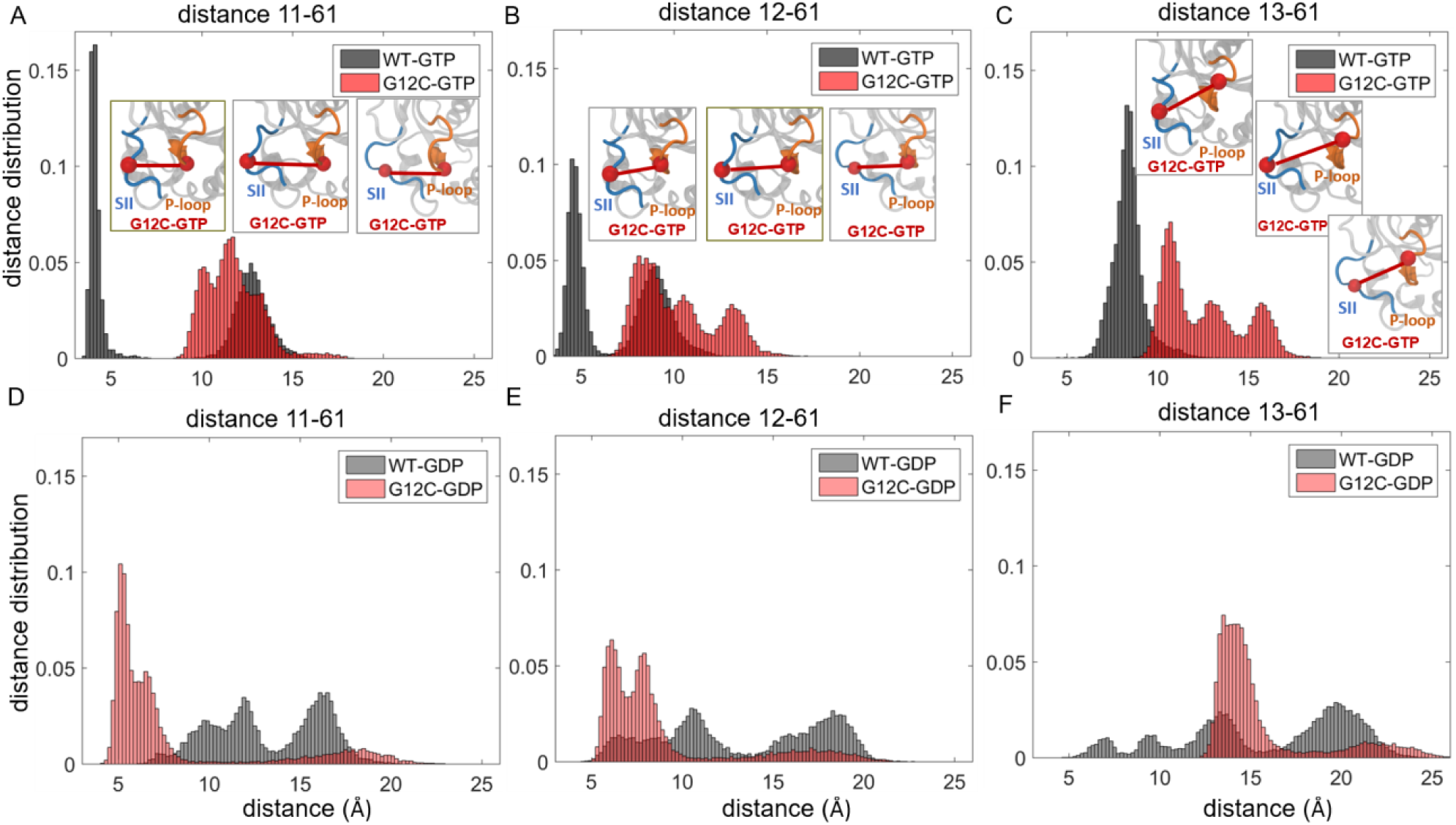
Distribution of distances *W*(*R*_*ij*_) between Cα-Cα atoms of residue pairs in the P loop-SII region. GTP-bound active K-Ras (black: WT, red: G12C mutant) (A) A11-Q61, (B) G12D-Q61, (C) G12D-Q61; GTP-bound active K-Ras (grey: WT, pink: G12C mutant) (D) A11-Q61, (E) G12D-Q61, (F) G12D-Q61.

In contrast to the widely dispersed distances in inactive form, *active K-Ras*^*WT*^ P-loop-SII distances show bimodal behavior with narrow ranges, which suggest two distinct conformations. While the P-loop-SII region can switch between the close and distant conformations in active K-Ras^WT^, in *all active studied mutations* it populates distant residue pair conformations within a wider range, including P-loop mutations at G12C (Fig 4A-C), G12V (Fig S2A-B), G13D (Fig S2C) as well as at Q61H (Fig S3). We have observed the same distribution pattern in K-Ras^G12D^ in our previous work^1^.

#### 2.1.5. In active K-Ras, mutations at P-loop cause α3 helix to move away from its neighbor, Loop7

In *inactive K-Ras*, the distances between α3 and Loop7 residues exhibit broad distributions, with two peaks in K-Ras^WT^ and one peak in mutant proteins K-Ras^G12C^, K-Ras^G12V^ and K-Ras^G13D^ (Fig 5, S4). However, upon activation, while these residue pair distances become less variable in K-Ras^WT^ with narrow distribution curves, they become more distant and variable in the mutants with multiple-peak distance distribution curves (Fig 4, S4).

**Figure 5.**
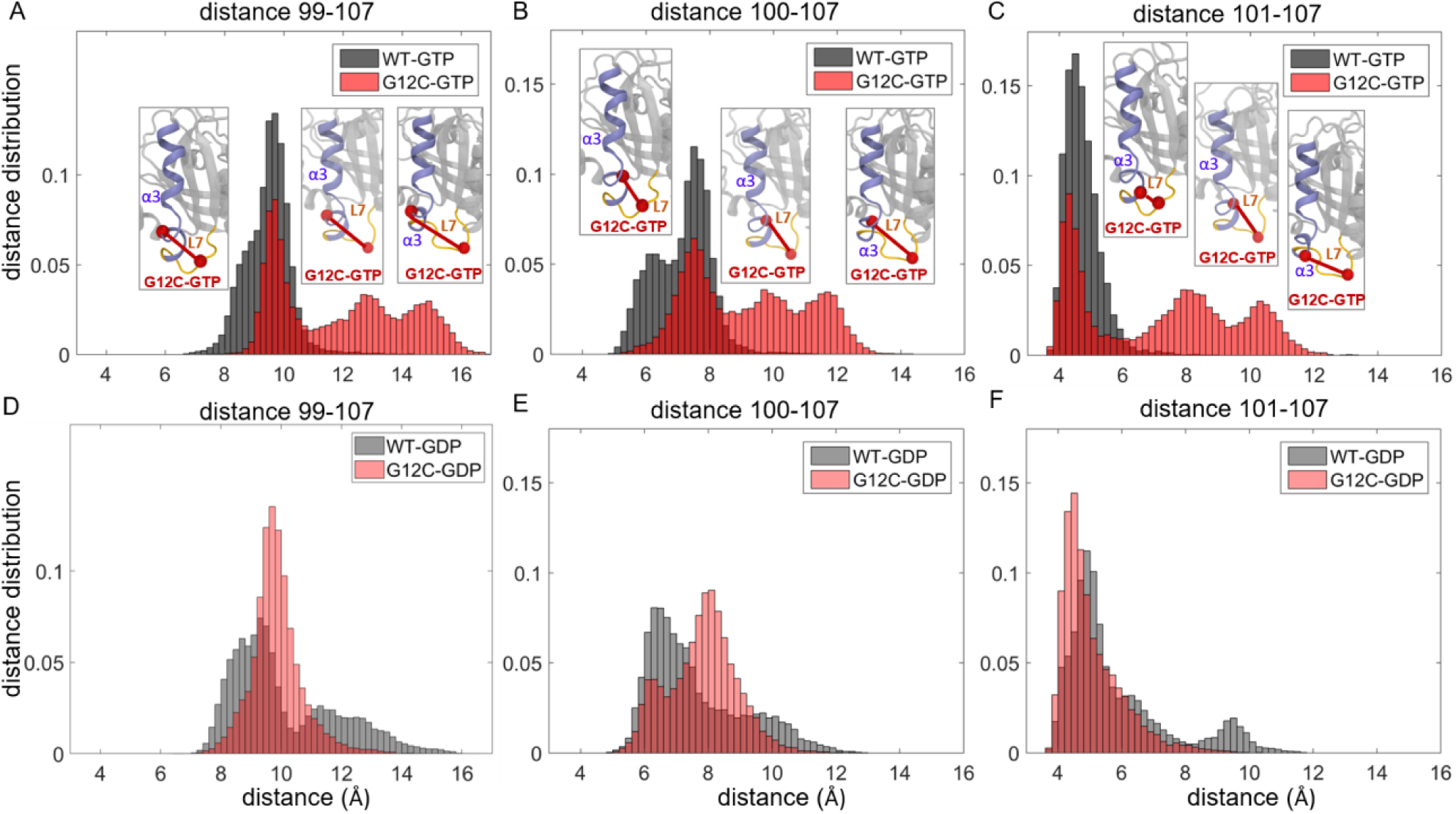
Distribution of distances *W*(*R*_*ij*_) between Cα-Cα atoms of residue pairs in the α3-Loop7 region. GTP-bound active K-Ras (black: WT, red: G12C mutant) (A) A11-Q61, (B) G12D-Q61, (C) G12D-Q61; GTP-bound active K-Ras (grey: WT, pink: G12C mutant) (D) A11-Q61, (E) G12D-Q61, (F) G12D-Q61.

#### 2.1.6. In active K-Ras, the G12C mutation causes the conformations of SII loop residues to become similar to those of inactive K-Ras^WT^

Distance distribution curves of the two residue pairs within the SII loop, T58-Q61 and Q61-64, show the same characteristics in both K-Ras^WT^-GDP (Fig S5C-D) and K-Ras^G12C^-GTP (Fig S5A-B). These curves have multiple peaks within the range of ∼5-10 Å that are related to multiple conformations of this region in K-Ras^WT^-GDP and K-Ras^G12C^-GTP. However, their single-peak narrow distribution curves in K-Ras^WT^-GTP implies the invariable conformation of the residue pairs in SII.

#### 2.1.7. In active K-Ras, all oncogenic mutations studied cause α1 and α5 helices to get stuck in their close conformation

In addition to the residue pairs above that move *away* from each other upon mutations as described above, we also observed regions that move *towards* each other, as shown in 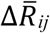 maps in Fig S6. Specifically, the negative 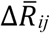 values indicate that the α1 and α5 helices get closer for all studied mutants of active K-Ras. Furthermore, their distance distribution curves each have a single peak (Fig S6A-D), indicating that the close conformation of the α1-α5 region becomes dominant upon mutations. In contrast, in wild-type protein, the two-peaked distance distribution curves of residue pairs (Fig S6A-D) suggests that they can obtain either a close or distant conformation. However, for inactive K-Ras, we did not observe such a conformational shift between the wild-type and mutant proteins (Fig S6E-H).

#### 2.1.8. In active K-Ras, all oncogenic mutations studied cause α2 helix of SII to move towards C-terminal of the α3 helix

In the inactive protein, the distances between D69 (α2, SII) and R102-V103 (α3) vary in the same range in both wild-type and mutant forms (Fig S7E-H). Conversely, in the active protein, the range of the distance values are not the same between the wild-type and mutant forms (Fig S7A-D). As evident from the distance distribution plots of the active K-Ras, D69 (α2, SII) and R102-V103 (α3) become closer in K-Ras^G12C^ followed by K-Ras^G12V^, K-Ras^Q61H^, K-Ras^G12D^ and K-Ras^G13D^.

### 2.2. DYNAMIC CHANGES IN K-RAS DUE TO ONCOGENIC MUTATIONS

#### 2.2.1. In active K-Ras, oncogenic mutations on the P-loop increase the flexibility of SII region

Comparing the fluctuation amplitudes of residues in active wild-type versus mutant K-Ras, we show that SI, Loop_β_2-_β_3, SII and α3-Loop7 regions are the flexible protein parts in all the P-loop mutant complexes G12C, G12V and G13D (Fig 6A-C). Moreover, we observed that the salt bridge between E63 (SII) and R68 (SII) in the active K-Ras^WT^ does not exist in the active P-loop mutants. In the absence of that bond, the flexibility of SII increases upon mutations in the P-loop and this region becomes the most fluctuating part of the mutant proteins (Fig 6A-C).

**Figure 6.**
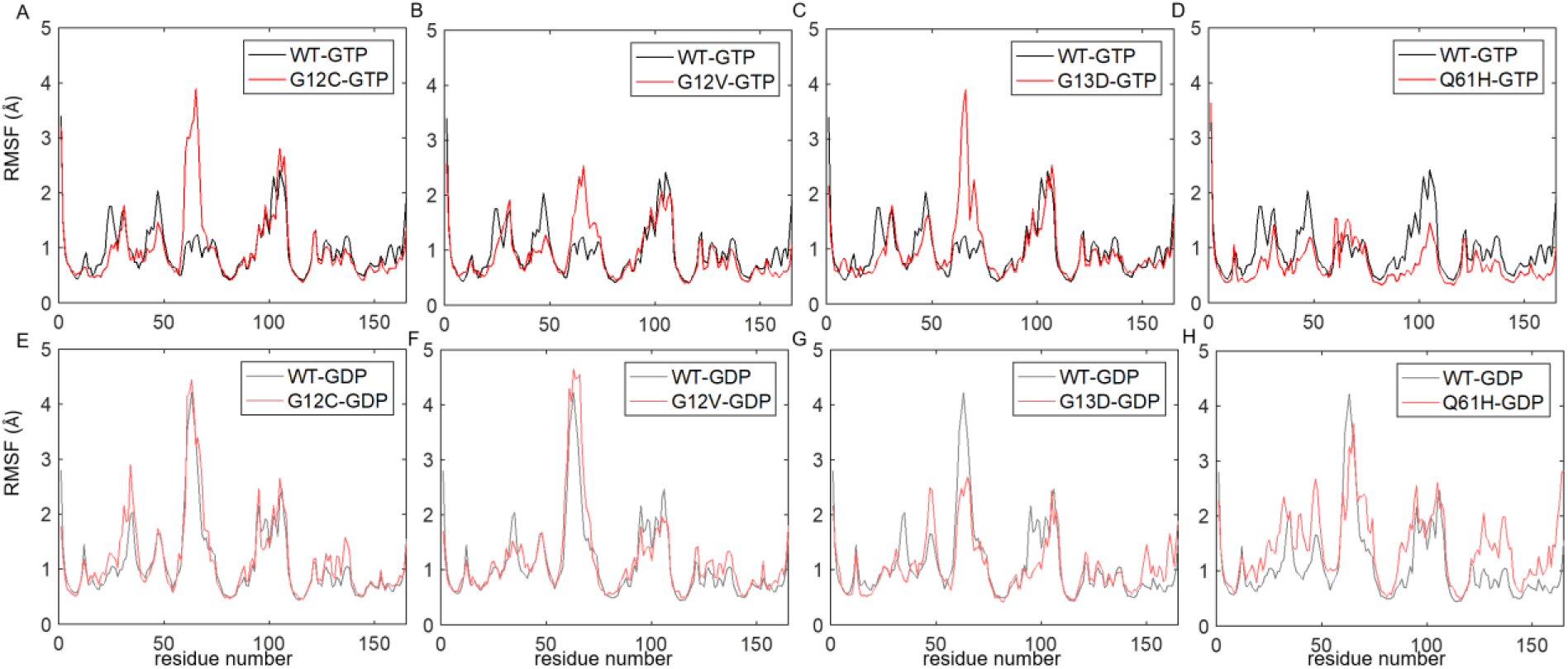
Changes in K-Ras dynamics due to oncogenic mutations. Y-axis shows the RMSF values of residues. GTP-bound active K-Ras (black: WT, red: mutant) (A) G12C-GTP (B) G12V-GTP (C) G13D-GTP (D) Q61H-GTP; GTP-bound active K-Ras (grey: WT, pink: mutant) (E) G12C-GDP (F) G12V-GDP (G) G13D-GDP (H) Q61H-GDP.

#### 2.2.2. In inactive K-Ras, G12C and G12V mutations do not affect the residue fluctuations

We compared the residue fluctuations of K-Ras^WT^-GDP with those of mutant K-Ras-GDP (Fig 6E-H). Root mean square fluctuation (RMSF) plots of K-Ras^G12C^-GDP and K-Ras^G12V^-GDP show that they have similar fluctuations to those of wild-type. Specifically, SI, SII and α3-Loop7 regions are the most flexible parts in both wild-type and mutant inactive proteins. Therefore, we conclude that G12C and G12V mutations do not change the flexibility of the inactive K-Ras.

#### 2.2.3. In inactive K-Ras, G13D mutation decreases the flexibility of SII (Fig 6G)

Comparison of RMSF plots of the inactive K-Ras complexes shows that SII residues fluctuate less in K-Ras^G13D^-GDP than in K-Ras^WT^-GDP. This indicates that the SII region of the GDP-bound K-Ras becomes less flexible due to the G13D mutations.

#### 2.2.4. The Q61 mutation alters protein flexibility differently than P-loop mutations (Fig 6D, H)

Although oncogenic mutations on P-loop increase SII fluctuations, the Q61H mutation on SII does not affect SII fluctuations in active K-Ras. Interestingly, Q61H mutation slightly decreases the fluctuations of other protein parts, indicating the rigid nature of K-Ras^Q61H^-GTP. Specifically, we observed that there is a salt bridge between residues E62 (SII) and K16 (α1). As a result of this bond, the amplitude of SII fluctuations in K-Ras^Q61H^-GTP are similar to those in K-Ras^WT^-GTP. However, Q61H mutation affects the flexibility of inactive protein in an opposite way. Most parts of the inactive protein become more flexible due to the Q61H mutation as indicated by the slightly higher RMSF fluctuation values in K-Ras^Q61H^-GDP.

### 2.3. CORRELATED RESIDUE PAIR MOTIONS IN K-RAS DYNAMICS

#### 2.3.1. Oncogenic K-Ras mutations do not disrupt correlated fluctuations of the residue pairs within the central β-sheet

In previous work, we have observed that the residue pair correlation maps of active (GTP-bound) and inactive (GDP-bound) K-Ras^WT^ show that fluctuations of the residues at the center of a six-stranded β-sheet are positively correlated (For details, see Vatansever *et al*^*1*^). Here, in residue pair correlation maps of the mutant proteins, we observe similar positive correlation patterns between the β-strand residues (Fig 7-8). Therefore, oncogenic mutations of K-Ras do not appear to affect the coupling of β-strand fluctuations.

**Figure 7.**
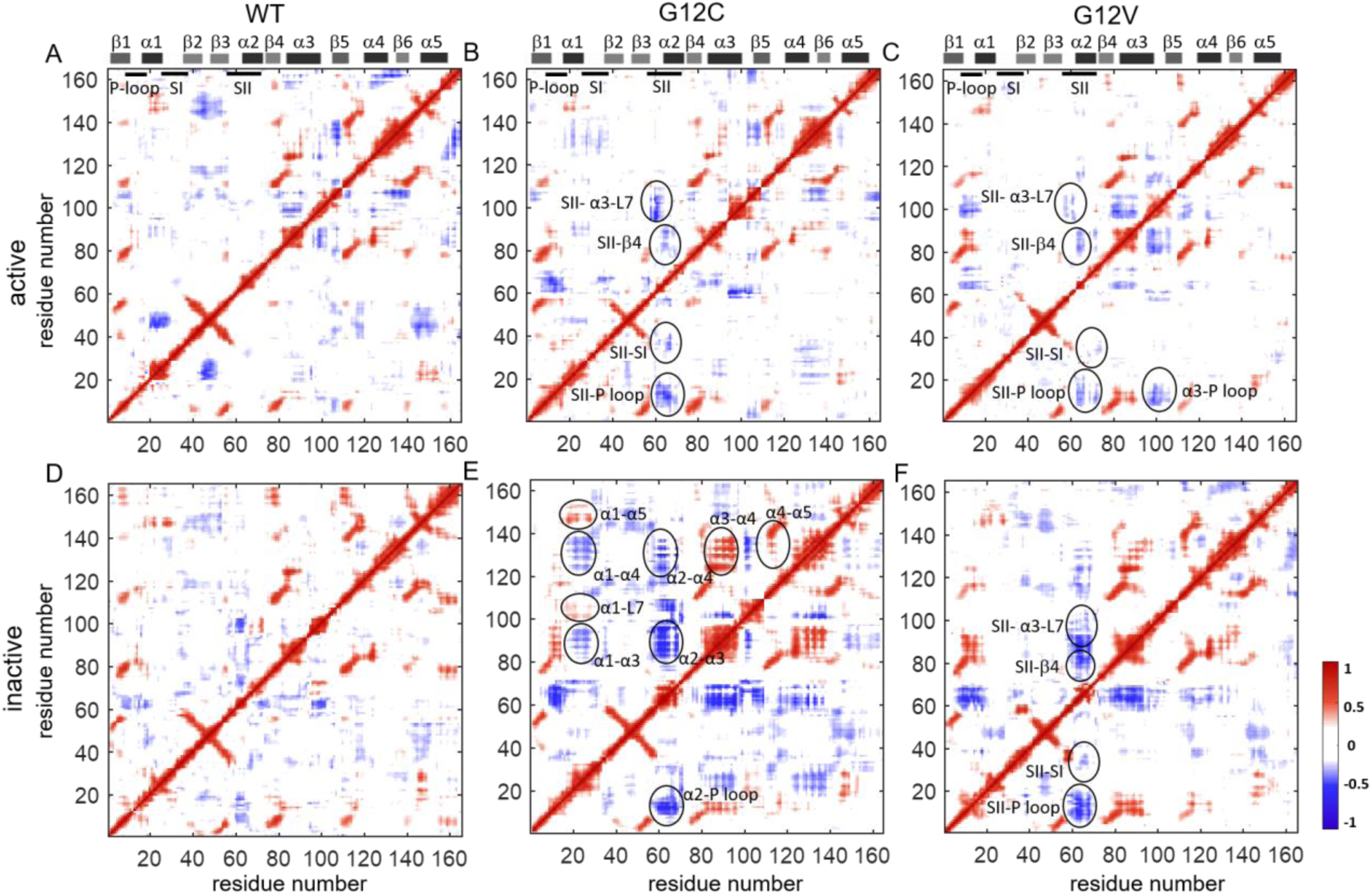
Correlated motions of residue pairs in both active and inactive K-Ras G12 mutants. Positive correlations are in red and negative correlations are in blue. Pairwise correlation coefficients are plot for (A) K-Ras^WT^-GTP, (B) K-Ras^G12C^-GTP, (C) K-Ras^G12V^-GTP, (D) K-Ras^WT^-GDP, (E) K-Ras^G12C^-GDP, (F) K-Ras^G12V^-GDP.

#### 2.3.2. Oncogenic mutations at residue G12 alter negatively correlated motions in K-Ras

From the pairwise correlation analysis, we observed that the negative residue pair correlations of K-Ras are G12 mutation-specific, as shown in Figure 7. Specifically, in the active wild-type protein, α1 and SI move in correlation with the β2-β3 regions; but this correlation is disturbed by the G12 mutations and new correlations occur. SII motions become strongly correlated with the P-loop, SI, β4 and α3-Loop7 regions upon G12C and G12V mutations. Additionally, α3 motions become correlated with the guanine nucleotide binding sites due to G12V mutation. For the inactive states, correlations between the SII motions and the other protein regions increase in the G12C and G12V mutant.

#### 2.3.3. In inactive K-Ras, G12C mutation significantly increases residue motion couplings

In the residue pair correlation analysis maps in Figure 7, inactive G12C mutant map displays the most striking changes, where motions of the α-helices become correlated with other protein regions (Fig 7E). Specifically, α1 motions are coupled to the Loop7 and α5 motions while they are negatively correlated with α3 and α4. Moreover, α2 motions are negatively correlated with the P-loop, α3 and α4. Furthermore, α4 moves in correlation with α3 and α5.

#### 2.3.4. In active K-Ras, G13D mutation weakens the correlations of SII while causing new correlations for β2-β3 loop (Fig 8A, C)

SII motions in G13D mutant show similar, albeit much weaker correlation patterns with those in WT, G12C and G12D complexes. However, β2-β3 loop motions become negatively correlated with the central β-sheet (β1-6) and positively correlated with SII. Additionally, α4 region moves in correlation with β1, the β2-β3 loop, β4, α3, β5 and α5.

**Figure 8.**
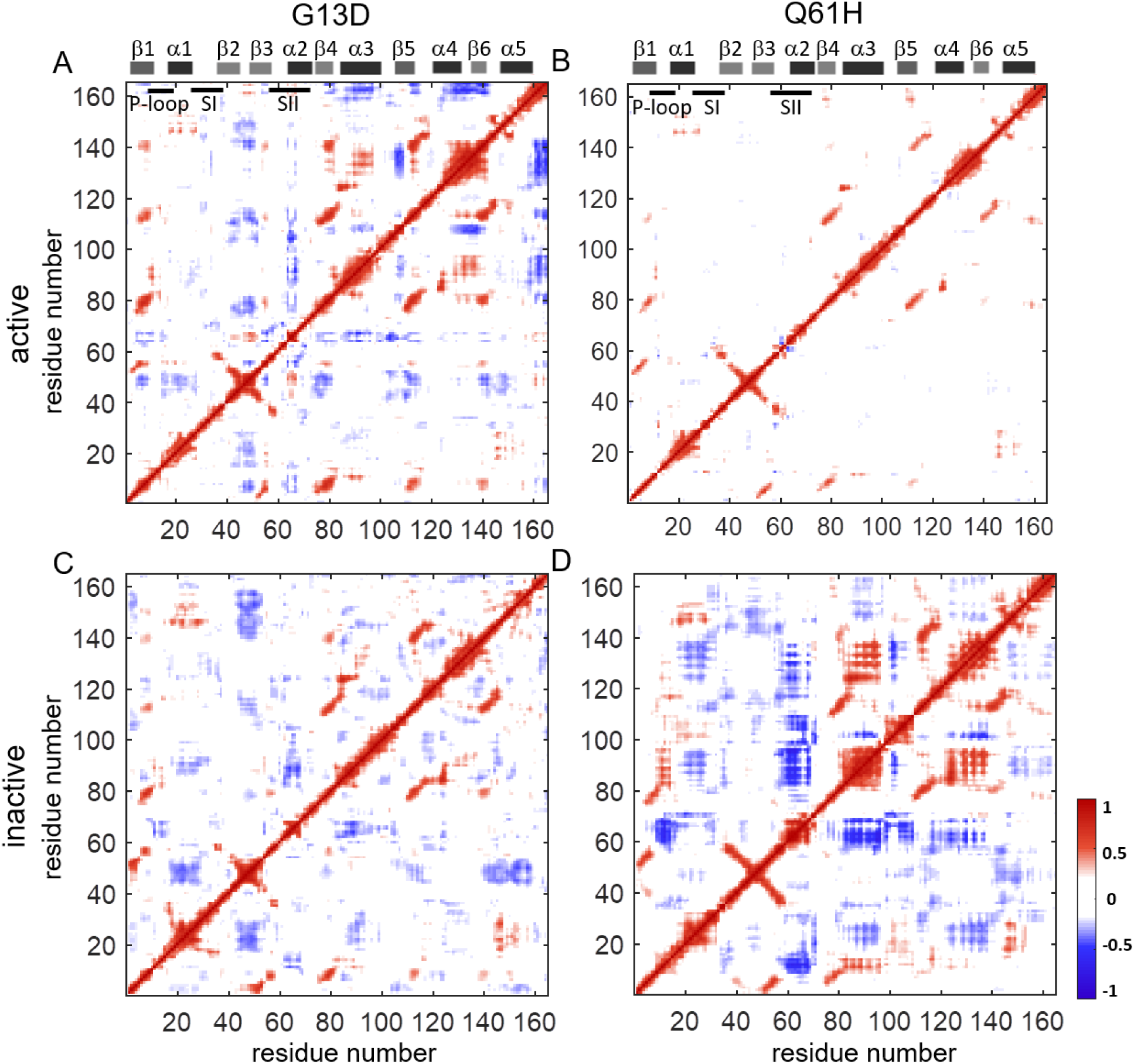
Correlated motions of residue pairs in both active and inactive G13D and Q61H mutants. Positive correlations are in red and negative correlations are in blue. Pairwise correlation coefficients are plot for (A) K-Ras^G13D^-GTP, (B) K-Ras^Q61H^-GTP, (C) K-Ras^G13D^-GDP, (D) K-Ras^Q61H^-GDP.

#### 2.3.5. Q61H mutation destroys correlated motions in K-Ras dynamics

The negatively correlated residue pair motions in K-Ras completely disappear in active Q61H complex, as shown in Figure 8B. On the other hand, Q61H mutation causes correlated motions of α1 the β2-β3 loop in the inactive protein (Fig 8D).

## 3. DISCUSSION

Oncogenic K-Ras mutants are high priority drug targets in cancer treatment. Hence, there have been considerable research efforts in directly targeting specific mutations or nucleotide-bound states (active or inactive) ^31^. Recently, nucleotide-specific inhibitors of K-Ras^G12C^ have shown promise^32^, which have fueled further efforts towards directly targeting other K-Ras mutants in active or inactive states. Such selective targeting efforts have led to comparative analyses^33^ that have improved our understanding of the mutation-specific effects at both clinical^34-36^ and molecular level^37-41^. For atomistic level comparative analyses, studies have utilized MD simulations to focus on the differences in the global dynamics of active and/or inactive K-Ras mutants ^22,23^. However, how different oncogenic mutations alter the *local* dynamics of K-Ras remains elusive. To address this question, in our previous work, we have studied the mechanisms by which oncogenic G12D mutation alters wild-type K-Ras conformations and dynamics and observed that it leads to changes specific to the bound nucleotide (GTP or GDP). Here, we build upon our previous study to understand whether the changes in nucleotide-bound K-Ras conformations and dynamics also show mutant-specific behavior. For this purpose, we investigated the changes in the conformational and dynamic behavior of active and inactive K-Ras caused by its most frequently observed oncogenic mutants other than G12D, which are G12C, G12V, G13D and Q61H. We analyzed the dynamics of each mutant protein in depth and compared them to those of the wild-type in both active and inactive forms. Our findings strongly sugggest that the intrinsic dynamic characteristics of the studied oncogenic K-Ras mutants are different than those of K-Ras^WT^. As other studies have also shown differences in K-Ras mutant oncoproteins in terms of transforming ability, GTP binding, anchorage-independent growth, and migration capacities^42^, our findings on differences in local dynamics can help better understand the mechanisms that underlie such mutant-specific differences in K-Ras biology. Collecting such information on local residue conformations and their dynamic features is also a critical step towards the design of mutant specific and potent small targeting molecules^43^. In summary, knowledge we have generated on mutant-specific local changes can help better understand the mechanisms that underlie the biological differences between K-Ras mutants, and further inform targeted drug design.

To evaluate mutation-specific changes in K-Ras conformations, we have used residue pair distance calculations. Figure 2 shows residue pair distance maps and Figures 3-5 show their distance distributions, which reveal significantly increased distances caused by oncogenic mutations. As shown in Figures 2-4, residues Q61-E62 (SII) move away from A11-G13 (P-loop) and D92(α3) in the active G12 mutant complexes. We also observed changes in the protein structure, where the salt bridge between E62 (SII) and K88 (α3) in K-Ras^WT^-GTP disappears upon mutations. Our integrated structural and conformational analysis revealed that G12 mutations disturb this salt bridge between SII and α3, leading SII to move away from its neighbors. The importance of the connections within this SII-α3 region for the K-Ras dynamics was emphasized in previous studies^44^. First, it was shown that the interactions between SII and α3 (i.e. M67-Q99/I100, F78/C80-I100) is specific to K-Ras by comparing it with the other isoforms –N-Ras and H-Ras. Then, this SII-α3 region was defined as an “allosteric switch” in active K-Ras based on the observation of a shift in relative SII-α3 conformations. Consistent with these studies, we identified specific effects of the studied oncogenic K-Ras mutations on structure and conformations of this region. Other studies have established that K-Ras mutations significantly reduce GAP-mediated GTP hydrolysis^45^, where SII region is the GAP-binding site. Hence, mutational changes in SII conformations relative to those of other regions may potentially disrupt GAP binding and thereby GAP-assisted GTPase activity.

Our distance analysis further reveals that all studied oncogenic mutations cause α1 and α5 helices to get stuck in their close conformation in active K-Ras (K-Ras-GTP). Previous studies have emphasized the importance of the interaction of the α5 helix with the lipidated hypervariable region (HVR) and membrane lipids for the K-Ras membrane orientation. These studies have used MD simulations with the HVR attached to the lipid membrane to understand the membrane orientation of the catalytic domain of an oncogenic K-Ras (i.e. G12V-KRAS^46,47^), but have not compared different mutant proteins or nucleotide states. Therefore, MD simulations of different membrane-bound oncogenic K-Ras proteins, as well as their comparative analyses are useful for further understanding the effects of oncogenic mutations on K-Ras dynamics.

To quantify how the studied oncogenic mutations change dynamic behavior of K-Ras, we used RMSF calculations, which showed higher fluctuation amplitudes for SII residues in active P-loop mutants as compared to wild-type (Figure 6A-C). We further observed that this change in protein flexibility is associated with other structural changes, including the disappearance of a salt bridge within the SII region (i.e., E63-R68). However, unlike the P-loop mutants, Q61H mutation does not alter SII fluctuations (Figure 7D), as another salt bridge forms between E62 (SII) and K16 (α1) in the active protein. This salt bridge disappears in inactive K-Ras^Q61H^-GDP, leading to increased SII fluctuations (Fig 5H). However, in terms of the flexibility of GDP-bound inactive protein complexes, only the G13D mutant differs from the wild-type. Notably, SII fluctuations only decrease in the G13D mutant, as depicted in the RMSF plot (Figure 6G). This finding is consistent with experimental studies^45^ that have shown that the GDP to GTP nucleotide exchange is similar for K-Ras^WT^ and all its mutants, but is faster in the G13D mutant. While this may contribute to the more aggressive biology of tumors with G13D mutation^45^, its combination with the decreased flexibility of the nucleotide-binding site SII may also provide a more accessible active site for small-molecules that bind to G13D mutant K-Ras.

We next investigated if and how the studied oncogenic mutations alter the fluctuations of correlated residue pairs, as these can be crucial in the regulation of protein dynamics, and thereby protein activity^48^. For this purpose, we calculated and compared the pairwise residue correlation coefficients of the mutant proteins, maps of which showed mutation-specific patterns (Fig 7-8). Specifically, in active G12 mutants G12C and G12V, pairwise residue fluctuations in SII become negatively correlated, which we have previously also observed in G12D^1^. We next chose residues representative of the SII region in these two mutants (Q61 in K-Ras^G12C^-GTP and S65 in K-Ras^G12V^-GTP), and studied in detail their correlations with other protein parts (Fig S7A-B). In K-Ras^G12C^-GTP, Q61(SII) shows negative correlations with residues 96-105 (α3-Loop7) and 10-17 (P-loop). As we discovered in our distance calculations, in G12C mutants SII also becomes distant from the α3-Loop7 and P-loop regions. Similarly, in K-Ras^G12V^-GTP, S65 (SII) becomes negatively correlated with and distant from residues S89 (α3) and 10-14 (P-loop). In summary, we have observed that in active K-Ras, G12 mutations cause SII to negatively correlate with and move away from α3-Loop7 and P-loop regions. Next, we studied the correlations of these residues in inactive K-Ras. We observed that in K-Ras^G12C^-GDP, Q61 (SII) shows strong negative correlations with C12 (P-loop) and H95 (α3). The distance distribution plots show that SII and P-loop populate distant conformations compared to K-Ras^WT^-GTP, while SII-α3 regions become closer. On the other hand, in K-Ras^G12V^-GDP, Q61 (SII) shows negative correlations with and becomes distant from the P-loop and α3. In the K-Ras^G13D^-GTP complex, Y64 (SII) exhibits negative correlations with C-terminal of the β3 (Fig S7C) while it moves away from the same region (Fig 2E). In active Q61H mutant, G12 fluctuates in negative correlation with SII region (Fig S7F) and moves away from the SII residues (Fig S3).

Combining coefficient calculations with the RMSF values, we observed that SII fluctuations become negatively correlated with other residue pair fluctuations when their amplitudes increase due to the mutations. Specifically, SII fluctuations increase in K-Ras^G12C^-GTP, K-Ras^G12V^-GTP and K-Ras^G13D^-GTP (Fig 6A-C) and become negatively correlated with the fluctuations of other residues (Fig 7B-C, 8A). We also observed the opposite of that relation in the K-Ras^Q61H^-GTP dynamics. SII fluctuations do not increase after Q61H mutation and they do not become correlated with the other protein parts.

In summary, we defined the unique dynamic behavior of oncogenic K-Ras mutants most frequently observed in cancer patients by utilizing long-time scale MD simulations. Comparative analysis of simulation data revealed common and distinct mechanisms of how oncogenic mutations affect K-Ras local conformations and dynamics. Associating the structural, conformational and dynamic alterations in active and inactive mutant proteins, our results provide a holistic understanding of effects of the mutations on local dynamics of K-Ras proteins. Such an understanding of the oncogenic mutant-specific dynamic characteristics of K-Ras proteins can inform the development of new direct inhibitor small molecules that selectively bind to active or inactive mutant K-Ras proteins.

## 4. METHODS

### 4.1. MD Simulations

We performed all-atom MD simulations of K-Ras^G12C^, K-Ras^G12V^, K-Ras^G13D^ and K-Ras^Q61H^ proteins bound to Mg^2+^GTP or Mg^2+^GDP. For construction of initial structures, we followed the analysis steps in our previous study (For details see Supplementary)^1^. Using NAMD 2.10 ^49^ with AMBER ff99SB ^50^ and general amber force fields (GAFF) ^51^, we performed MD simulations of each mutant protein again following the steps from our previous study ^52^, details of which we present in Supplementary. We ran 1 microsecond MD simulations of each complex in replicate and analyzed the last 900ns of each simulation. We recorded the atomic coordinates 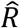 from the simulation trajectories in every 10ps and aligned them to the first frame using VMD software 1.9.2 ^53^. To identify salt bridge formations within the protein residues, we used Salt Bridges Plugin, Version 1.1, of VMD. We visualized the trajectories with VMD.

### 4.2. Residue pair distance calculations

We quantified the oncogenic mutation effects on residue pair distances using a computational approach described in our previous work^1^. In detail, based on the Gaussian network model (GNM)^24-27^, we assumed the maximum Cα-Cα distance separation between two contacting residues as ∼7.2Å^28^, and thereby defined a volume V with a radius of r1 ∼7.2 Å as the ‘first coordination shell’. In addition to the residue contacts within the ‘first coordination shell’, the contribution of non-bonded interactions to higher-order coordination shells can be important^29,30^. Therefore, we defined the ‘second coordination shell’ at twice the volume of the first, with a radius of ∼9.1 Å and included the residue pairs within their ‘second coordination shell’ in K-Ras structures in our residue pair distance analysis^30^. Briefly, we computed the Cα-Cα distances of residue pairs (*i, j)* in wild-type K-Ras (the reference state) and in mutant K-Ras complexes (the final state) (For details see Supplementary). For every residue pair *(i, j)* within the second coordination shell (radius of ∼9.1 Å) we calculated its time-averaged distance in K-Ras^WT^ 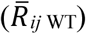, in mutant K-Ras 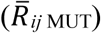 and the difference 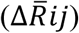, where 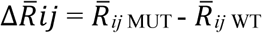 The magnitude of the difference value is the degree of conformational change caused by the mutation. The left column of Figure 1 presents residue pair distance maps in K-Ras mutants, where positive 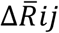 values suggest that a residue *i* moves away from residue *j* and negative values suggest that the residues get closer.

To quantify the changes in local volumes due to the mutations, we calculated the average of all 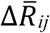 values using the same approach in our previous study^1^. For all residues *j* in the second coordination shell of residue *i*, we computed averaged 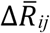 values for each residue *i* based on the formula of 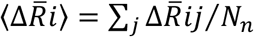, where *N*_*n*_ is the total number residues (*j*) in the second coordination shell of residue *i* (For details see Fig S8). The resulting 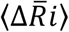 values provide a scale for the change in volume around each residue (*i*) upon mutation.

### 4.3. Distance distributions between residue pairs

We first calculated the pairwise distance, *R*_*ij*_ for each residue pair *(i, j)*, and then the normalized distribution of these distances, *W*(*R*_*ij*_), as we previously described^52^. Briefly, for every time point *t* of the MD simulation, *R*_*ij*_*(t)* simply corresponds to 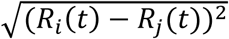. Next, to calculate *W*(*R*_*ij*_), we divided the range of *R*_*ij*_*(t)* across the time course of the simulation, (max{*R*_*ij*_*(t*_*1*_*), R*_*ij*_*(t*_*2*_),..*}-* min*{R*_*ij*_ *(t*_*1*_*), R*_*ij*_*(t*_*2*_),..*})*, into small bins of width 0.2 and counted the number of observed distances in each bin using the Histogram function of the MATLAB tool from MathWorks (Natick, MA). For example, for *W(R*_*ij*_) values of residue pair (61, 92), see y-axes of Figure 3.

### 4.4. Residue pair correlation calculations

We calculated the correlation coefficients of residue pair fluctuations to identify those whose fluctuations were coupled during protein motions. *C*_*ij*_ is the correlation coefficient value of a residue pair (*ij*) that varies from −1 to 1. For a perfectly positively correlated residue pair, C*ij* equals to 1, and to −1 for a perfectly negatively correlated residue pair. However, it equals to 0 if the residue pair fluctuations are not correlated. We computed the *C*_*ij*_ values for every residue pair in each K-Ras mutant using the algorithm in our previous study ^52^.

## 5. ACKNOWLEDGEMENTS

ZHG acknowledges funding from the LUNGevity Foundation and start-up funds from the Department of Genetics and Genomics and the Icahn Institute for Data Science and Genomic Technology. We would also like to thank Dr. Myvizhi Esai Selvan for help with one of the figures.

